# Prior Experience Alters the Appearance of Blurry Object Borders

**DOI:** 10.1101/701995

**Authors:** Diana C. Perez, Sarah M. Cook, Mary A. Peterson

## Abstract

Object memories activated by borders serve as priors for figure assignment: figures are more likely to be perceived on the side of a border where a well-known object is sketched. Do object memories also affect the *appearance* of object borders? Memories represent past experience with objects; memories of well-known objects include many with sharp borders because they are often fixated. We investigated whether object memories affect appearance by testing whether blurry borders appear sharper when they are contours of well-known objects versus matched novel objects. Participants viewed blurry versions of one familiar and one novel stimulus simultaneously for 180ms; then made comparative (Exp. 1) or equality judgments regarding perceived blur (Exps. 2-4). For equivalent levels of blur, the borders of well-known objects appeared sharper than those of novel objects. These results extend evidence for the influence of past experience to object *appearance*, consistent with dynamic interactive models of perception.

---

Studies of vision demonstrate that bottom-up processes are insufficient for understanding how the brain constructs our percept of the external world; instead, higher factors such as attention, expectation, and memory influence visual processing^1–5^. While attention facilitates processing of visual stimuli by granting priority to the more relevant aspects of the scene, memory and expectation constrain possible interpretations of visual input and predict what is likely to occur in the sensory environment; all result in improved perceptual performance^3, 6–8^.

In addition to behavioural effects, Carrasco and colleagues showed that spatial attention can increase the subjective *appearance* of contrast^9^. Their participants viewed two Gabor patches shown to the left and right of fixation; their task was to report the orientation of the higher contrast patch. Before the Gabor patches were shown, participants were cued to allocate attention to either the left or right location. They found that the stimulus in the location to which attention was allocated appeared higher in contrast than the unattended stimulus. Additional experiments extended the influence of attention on stimulus appearance to other attributes, such as spatial resolution^10^, spatial frequency and gap size^11^, colour saturation^12^, and motion coherence^13^.

The results described above raise the question of whether other high-level factors, such as expectation and prior experience, can modulate stimulus appearance, in particular the perceived sharpness of blurry object borders^14–15^. Like contrast and spatial frequency, blur is a fundamental feature in vision. Blur is present in varying degrees due to changes in the environment (distance or depth) or the observer (pupil diameter or refractive errors), yet the visual system compensates for blur to allow us to see the world ‘in focus’^16^. Blur detection thresholds have been shown to be unaffected by manipulations of attention via cognitive load^17^, but a previous study suggests that the appearance of blur is affected by past experience^15^: In order to match the apparent blur of a real word, a comparison pseudo-word had to be sharper. The stimuli in that experiment were displayed indefinitely; hence, observers could have adjusted their focus differently when looking at real versus pseudo-words, in which case the results would indicate that prior experience alters behaviour (which leads to a change in perceived blur) but would not indicate that prior experience directly alters appearance. We report six experiments investigating whether the integration of memory representations with sensory input can sharpen the appearance of blurry borders of a well-known object versus a matched novel object when both objects were briefly presented simultaneously, eliminating the possibility of differential accommodation for the two types of stimuli.

A well-known object is one that has been encountered often in daily life (henceforth, referred to as a *familiar* object). Therefore, memories of familiar objects probably span a range of blur levels. Many of these memories are likely to be sharp given that people fixate on objects with which they interact^18^. Thus, if percepts are generated by combining bottom-up input with higher-level object memories activated by a stimulus, then for a given level of blur, the borders of familiar objects should appear sharper than those of novel objects. We tested this hypothesis and also examined whether predictions generated from a word prime can sharpen the perception of the blurry borders of a well-known object. The presentation of a prime typically improves performance, most likely because the word activates a target-related concept before the stimulus^19–21^. We used a masked prime to attempt to manipulate expectations before the silhouettes appeared.

## Results

### Experiment 1 – Comparative Judgements

In Experiment 1A we presented two objects, one familiar and one novel. The novel object was matched to the familiar object by rearranging the parts of the familiar object into a novel configuration (Fig. 1A). The objects were blurred using a 2-D Gaussian smoothing filter. The standard deviation (σ) of the smoothing filter was manipulated from σ = 3 - 11, thereby manipulating blur over a range of 9 standard deviations that we refer to as blur levels (Fig 2; see Methods). In half the trials, the familiar object was the *Standard*, set at blur level of 7 and the novel object was the *Test*, presented at blur levels ranging 3 (least blurry) to 11 (blurriest); in the other half of the trials, these roles were reversed. Masked word primes preceded the stimulus presentation. These primes denoted either the familiar object, an unrelated object, or had no semantic value (a string of x’s; Fig. 3 shows a sample trial sequence). Participants were instructed to report whether the object on the left or on the right appeared blurrier.

**Figure 1.**
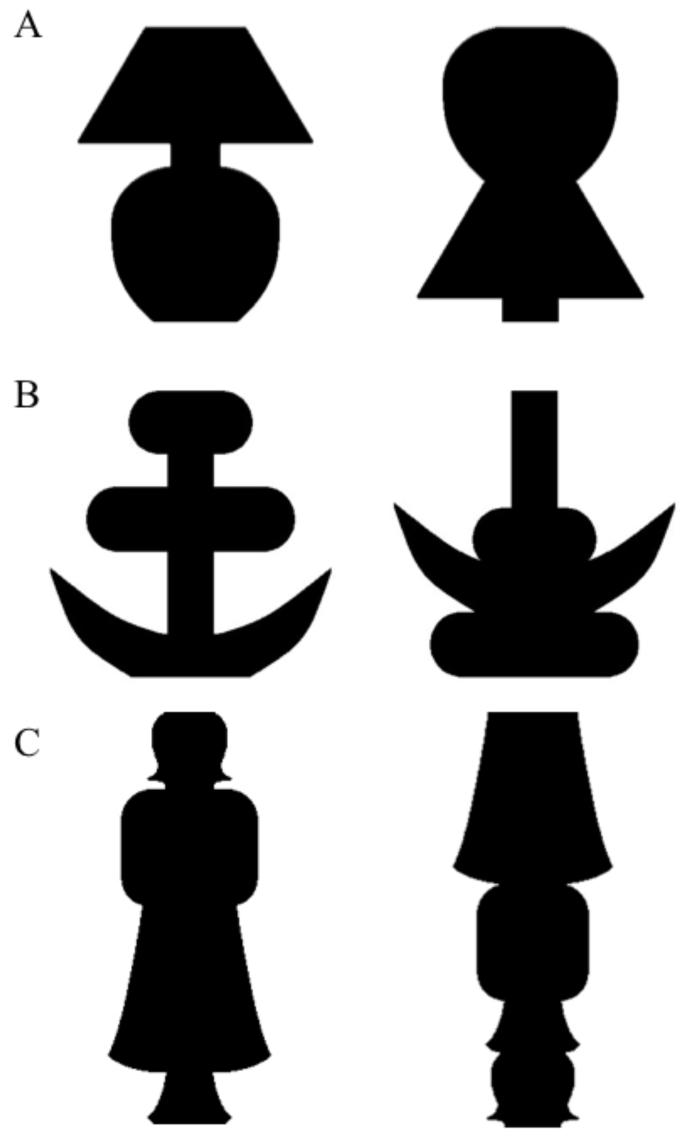
Stimuli used in Experiments 1-4. The familiar object is on the left and the novel object on the right. The novel stimulus was an object created by rearranging the parts of the familiar object into a different, novel, configuration, where parts were delimited by successive minima of curvature. The two types of objects appeared equally often to the left and right of fixation. The stimuli are shown here without blur. A) Stimuli used in Experiments 1-3; the familiar object is a table lamp. B) Stimuli used in Experiment 4A; the familiar object is an anchor. C) Stimuli used in Experiment 4B; the familiar object is a standing woman.

**Figure 2.**
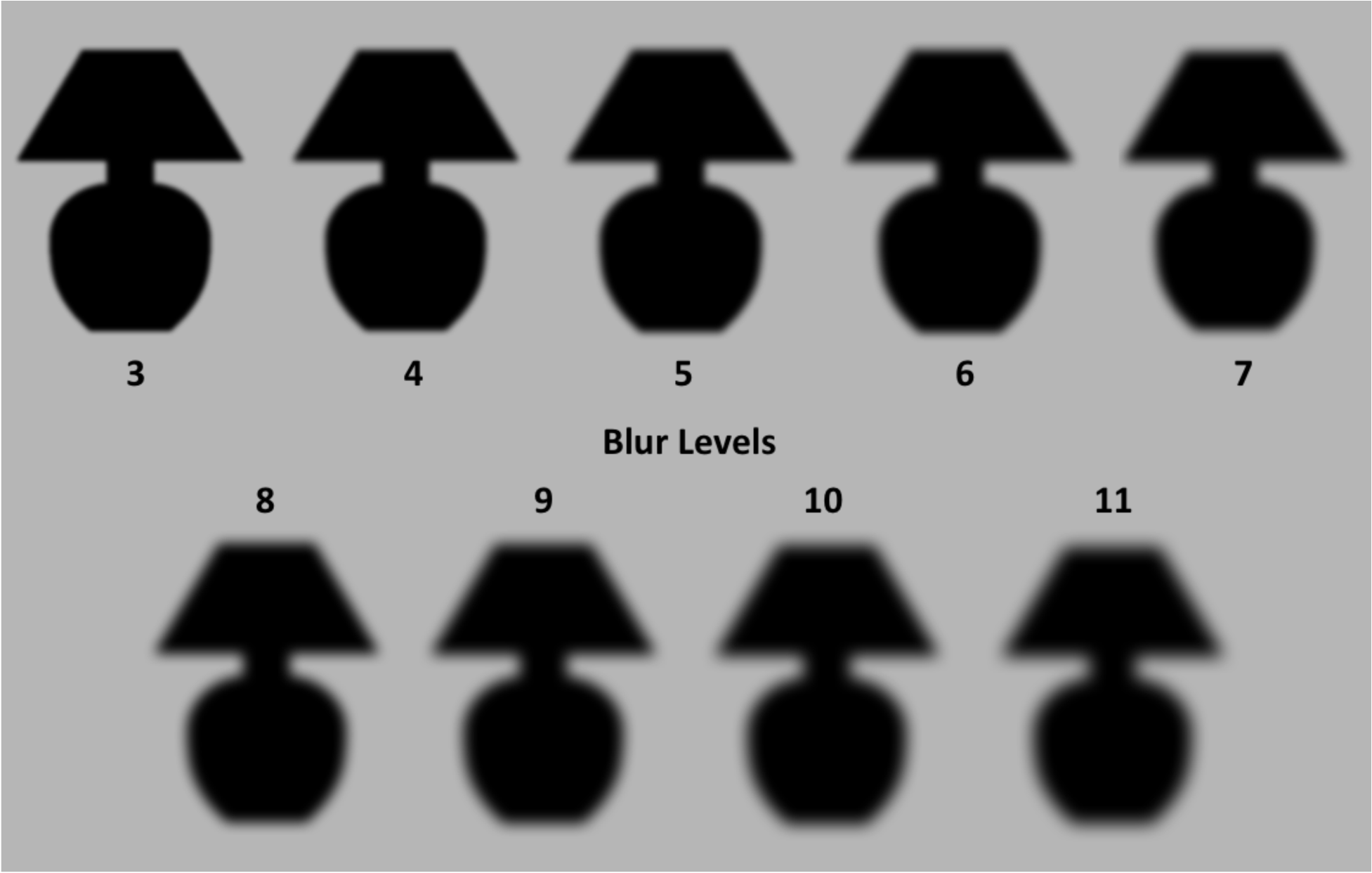
Blur levels for Lamp stimulus. Depiction of *Test* blur levels used for all experiments applied to the familiar stimulus used in Experiments 1-3, the silhouette of a lamp. The blur level refers to the standard deviation of the Gaussian distribution that gave its shape to the smoothing kernel used to blur the stimuli (see Methods for more explication). The blur level of the *Standard* stimulus was always 7.

**Figure 3.**
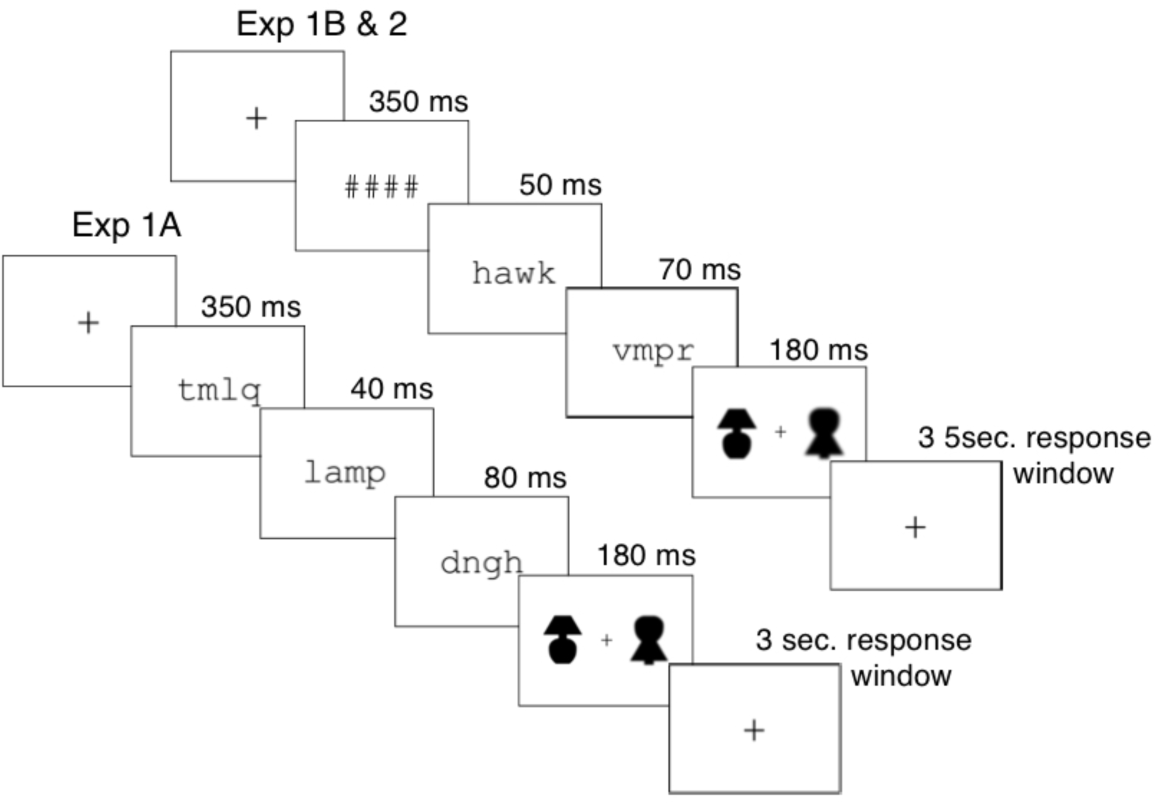
Sample trials for Experiments 1A (lower stream), and Experiments 1B and 2 (upper stream) Experiments 1A, and 1B and 2 used the same word primes, post-masks and stimuli, but differed in timing and pre-masks. In the sample trial sequence for Experiment 1A, the *Standard* (at blur level 7) was the novel object presented to the right of fixation, and the *Test* was the familiar object, illustrated here at blur level 3. In the sample trial sequence for Experiments 1B and 2, the *Standard* was the familiar object to the left of fixation, and the *Test* was the novel object, illustrated here at blur level 11. The word primes in this figure are enlarged for visibility.

As expected if object memories sharpen the appearance of blurry borders, the point at which *Test* objects appear equal to *Standard* objects was shifted toward the blurrier end of the continuum for familiar compared to novel *Test* objects, indicating that for a given level of blur, the borders of the familiar object appear sharper than those of the novel object. Fig. 4A shows the average proportion of trials on which participants reported the *Test* object was blurrier as a function of the *Test* object blur level. A logistic function fit to the averages yielded the point of subjective equality (PSE). Fig. 4B shows the PSE for each of the priming conditions and each stimulus condition. In Experiment 1A a blurrier familiar *Test* object (*PSE* 7.59, *SE* = 0.091) was subjectively equal to a *Standard* novel stimulus, and a sharper novel *Test* object (*PSE* 6.42, *SE* = 0.069) was subjectively equal to a *Standard* familiar stimulus. This difference was expected if perception results from the integration of the current sensory information with long-term memory (LTM) representations of objects that on average are sharper than the blurry versions of familiar objects presented in the experiment. Averaging over the PSE differences found in the two *Test* object conditions, the results show that the familiar object appeared sharper than the matched novel object by 0.59 of a blur step in Experiment 1A. A 2 × 3 ANOVA (with the factors of *Test* Object type: familiar vs. novel, and Prime type: identity, unrelated, and control), showed a statistically significant main effect of *Test* object type, *F*_1,13_ = 70.9, *p* < .00001, η_p_2 = 0.85. There was no effect of Prime type, *F*_2, 26_ = 1.57, *p* = 0.23, η_p_^2^ = 0.11; nor was there an interaction between Prime type and *Test* Object type, *F*_2, 26_ = 0.14, *p* = .87, η_p_^2^ = 0.010.

**Figure 4.**
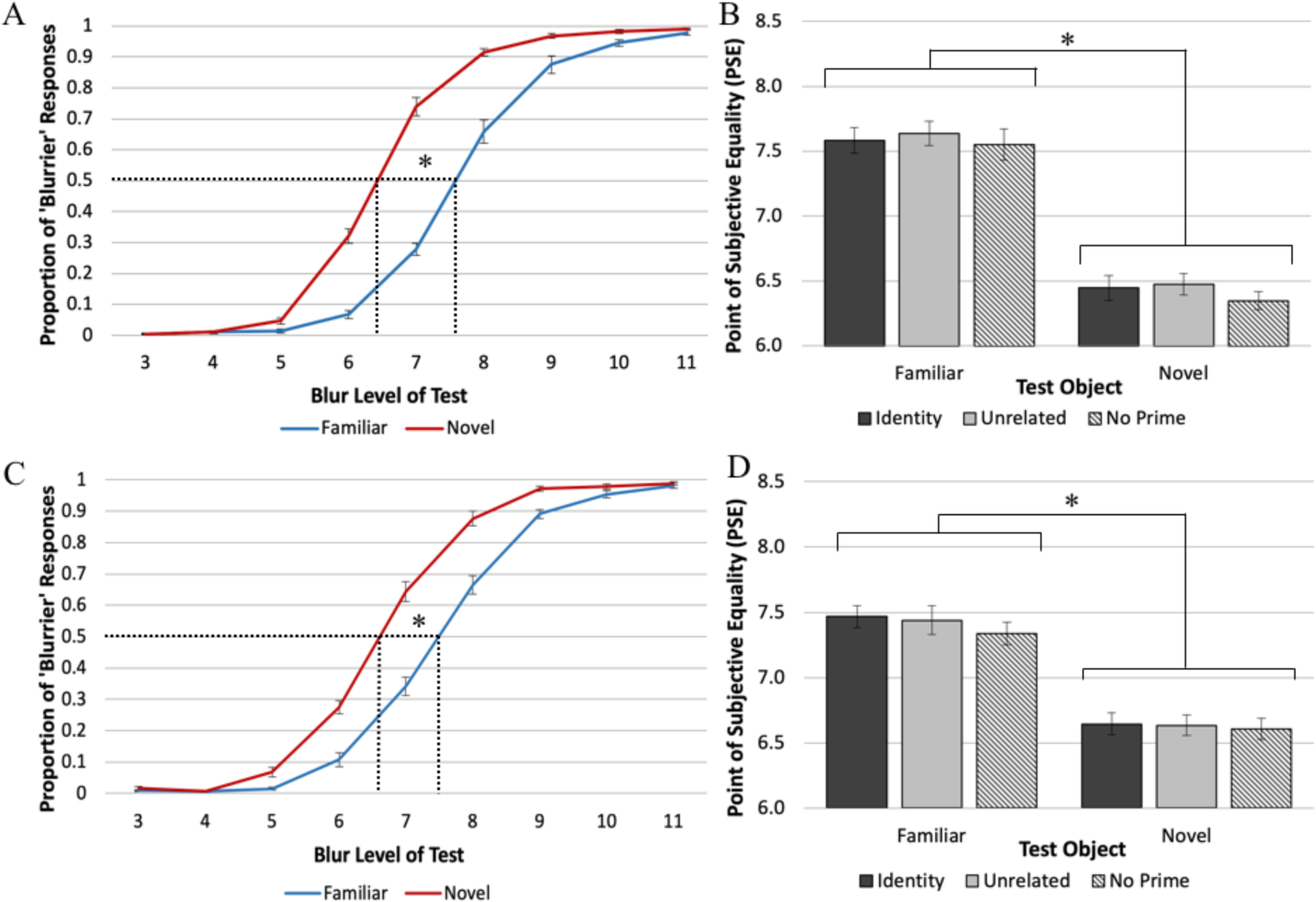
Results of Experiments 1A and 1B. A & C) The mean proportion of trials on which that the *Test* object was perceived as blurrier than the *Standard* (blur level 7) as a function of the blur level of the *Test* object for Exp. 1A (A) and Exp. 1B (C). The dotted lines indicate the PSEs. For familiar *Test* objects a blurrier stimulus was subjectively equal to a *Standard* novel stimulus and for novel *Test* objects a sharper stimulus was subjectively equal to a *Standard* familiar stimulus. This is consistent with the hypothesis that for a given level of blur, familiar objects appear sharper than novel objects. B & D). The PSE values for familiar and novel *Test* objects as a function of Prime type. No effects of Prime type were obtained. Errors bars represent the Standard Error.

We hypothesized that the absence of a priming effect might occur because the masks interfered with the effectiveness of the word prime. Accordingly, we changed the masking conditions in Experiment 1B, such that the pre- and post-masks were shorter in duration, whereas the prime was longer. To further reduce the interference of the pre-mask with the prime, on each trial, we substituted a string of pound signs (“####”) for the random string of consonants used in Experiment 1A.

Fig. 4C shows the proportion of ‘Blurrier’ responses as a function of *Test* object blur level for Experiment 1B. Replicating Experiment 1A, the curve-fitting data showed that blurrier familiar *Test* objects appeared subjectively equal to *Standard* novel objects (*PSE* = blur of 7.41, *SE* = 0.068) and sharper novel *Test* objects appeared equal to *Standard* familiar objects (*PSE* = blur of 6.63, *SE* = 0.060; Fig 3D). Thus, overall familiar objects appeared sharper by a blur step = 0.39. An ANOVA showed a statistically significant main effect of *Test* Object type, *F*_1,13_ = 46.5, *p* < .00001, η_p_2 = 0.78. Again, we observed no main effect of Prime type, *F*_2, 26_ = 0.56, *p* = 0.58, η_p2_ = 0.041, and no significant interaction between Prime type and *Test* Object type, *F*_2, 26_ = 0.30, *p* = 0.75, η_p2_ = 0.022.

The lack of priming effects may be due to the low predictive relationship of the identity prime to the stimuli: Both the familiar and the novel stimulus were shown on each trial; subjects were unaware of which was the *Test* and which was the *Standard*. The prime conditions referred only to the *Test* object; hence, the prime denoted the familiar *Test* stimulus on only 16.7% of trials; 33.3% of the trials were unprimed (control) trials, and 50% of the trials were preceded by a prime that was unrelated to the familiar object (see Methods). This low predictive relationship probably reduced the likelihood of observing a priming effect. It is clear that even for familiar *Test* objects, the primes had no effect.

In order to verify that we manipulated familiarity effectively, participants were asked to identify the two objects that were presented during the experiment on a post-experiment questionnaire. In Experiment 1A, 76% of participants named the familiar object as a lamp. The most common interpretation for the novel object was ‘ghost,’ with 32% agreement. In Experiment 1B, 66% of participants identified the familiar object. The most common interpretation for the novel object was again ‘ghost,’ with 25% agreement. Thus, participants were more likely to perceive one of the stimuli as denoting a familiar object than the other.

### Experiment 2 – Equality Judgements

In Experiment 2, participants were instructed to report whether the two objects presented on each trial were the same or different levels of blur. We recorded equality judgments to allay the potential response bias criticism that participants may have simply chosen the novel object when they were uncertain regarding which object was blurrier^22, 23^. We averaged the proportion of ‘Same’ responses for each *Test* blur level (Fig. 5A). We then fit these averages to a normal Gaussian distribution to obtain a centroid value (*b*), which corresponds to the location of the peak of the distribution (where the probability of a ‘Same’ response is highest). Fig. 5B shows the centroid value for each *Test* Object type and each Prime type in Experiment 2. Replicating the effect of Experiment 1, this time assessed by equality judgements rather than comparative judgements, the familiar object was perceived as sharper than the novel object (*b_familiar_* = blur level of 8.02, *SE* = 0.085; *b_novel_ =* blur level of 7.56, *SE* = 0.078). The ANOVA showed a significant main effect of *Test* Object type, *F*_1,22_ = 34.1, *p* < .0001, η_p_^2^ = 0.61. These results support the hypothesis that, for familiar objects, perceived sharpness is a joint function of the current stimulus and LTM representations of familiar objects that are on average sharper than the *Standard* stimulus in the current experiment. No such LTM representations exist for the novel object.

**Figure 5.**
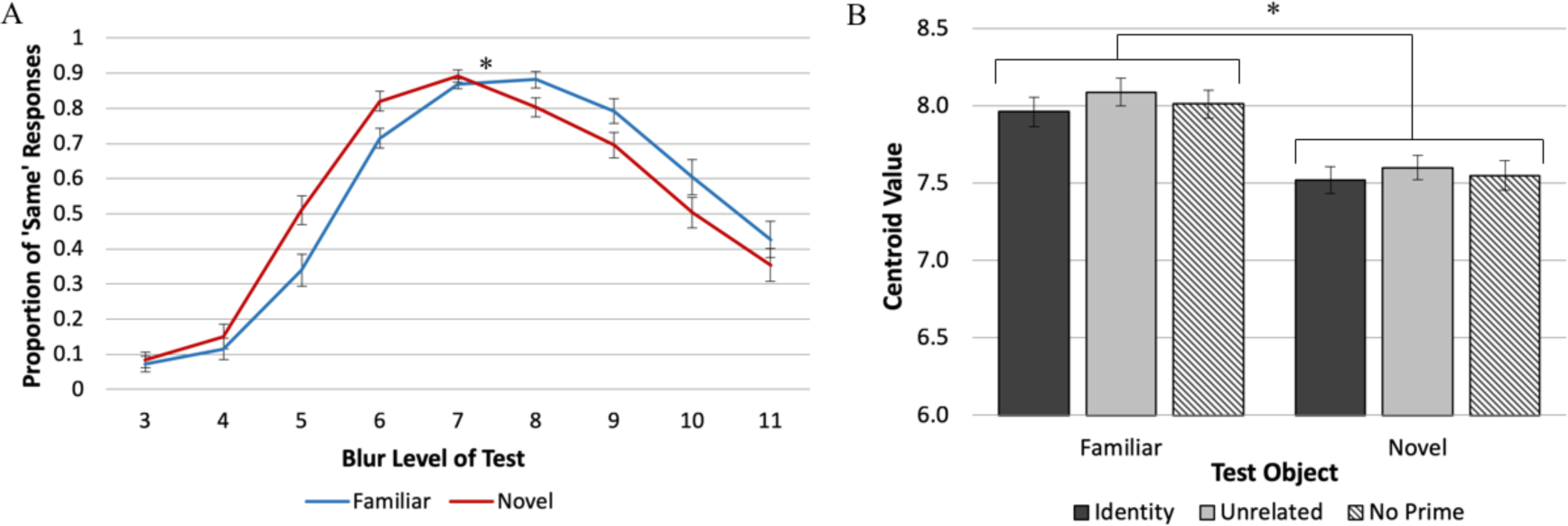
Results of Experiment 2. A) The mean proportion of ‘Same’ responses as a function of the blur level of the *Test* object. B) Centroid value estimates separated by *Test* object type and Prime type. The centroid values show that for familiar *Test* objects a blurrier stimulus was subjectively equal to a *Standard* novel stimulus of blur level 7. These results, obtained using equality judgments, are consistent with the hypothesis that for a given level of blur, familiar objects appear sharper than novel objects. Errors bars represent the Standard Error.

We also observed a main effect of Prime type, *F*_2,44_ = 3.65, *p* < 0.05 η_p_^2^ = 0.14: The centroid values for both familiar and novel *Test* objects were significantly higher when they were preceded by an unrelated prime (*b* = blur level of 7.85, *SE* = 0.068) rather than an identity prime (*b* = blur level of 7.74, *SE* = 0.078; p < .02). It is difficult to interpret this main effect because neither “unrelated” nor “identity” primes had any relationship to the novel object; they were dummy primes from the same categories as the primes for the familiar object – an artificial object in the identity condition (i.e., “pole”) and a natural object in the unrelated condition (i.e., “hawk) – yet the effect was present. Moreover, the mean centroids in the “no prime” condition did not differ significantly from those in the other two priming conditions, *p* > 0.28 for the identity prime and *p* > 0.13 for the unrelated prime, so the results can’t stem from a systematic response to the prime category (e.g., systematically inhibiting expectations based on artificial primes but not natural primes). Finally, Prime type and *Test* Object type did not interact, *F*_2, 44_ = 0.15, *p* = 0.87, η_p_^2^ = 0.007, as would be expected if the identity prime induced an expectation that modulated the perceived blur of the familiar object but not the novel object.

To estimate the perceived increase in sharpness due to familiarity, we divided the difference between the centroids for the familiar versus novel objects by two to account for the fact that any difference observed assays the perceived sharpening of the familiar object both when it was the *Test* and when it was the *Standard*, as in Experiments 1A and 1B. Using this method, the familiarity-dependent sharpening step was estimated as 0.23 (SE = 0.040) of a blur step, smaller than in Experiments 1A and 1B. We attribute this difference to evidence indicating that equality judgments are harder than comparative judgments and responses are more variable^24–26^. Nevertheless, the estimated sharpening step in Experiment 2 was statistically greater than 0, *t*_22_ = 5.84, *p* < 0.00001, *Cohen’s d* = 1.22.

In Experiment 2, the centroid for both the familiar and novel *Test* objects corresponded to a blurrier object than the *Standard*. This centroid shift may result from the asymmetric application of two criteria for ‘Different’ — “different because sharper” and “different because blurrier.” As Fig. 5A clearly shows, there are more ‘Same’ responses when the blur of the *Test* object is greater than that of the *Standard* object. This asymmetry likely arises from a known nonlinearity in responses to blur wherein increments in blur level are harder to discriminate for increasingly blurry stimuli^27, 28^. Asymmetric use of two criteria in equality judgments is not uncommon^24^. Note, however, that responses were affected by this asymmetry when both novel and familiar objects were the *Test* object. Therefore, the asymmetric use of ‘Different’ criteria cannot account for the results showing that for a given level of blur, familiar objects are perceived as sharper than novel objects.

Most participants in Experiment 2 identified the familiar object as a lamp with 59% agreement. The most common interpretation for the novel object was again ‘ghost,’ with 21% agreement. Thus, participants were more likely to perceive one of the stimuli as denoting a well-known “familiar” object than the other.

### Experiment 3 – Equality Judgements without Priming

In Experiment 3, participants again reported whether the two objects presented on each trial were the same or different levels or blur. Because of the absence of systematic effects of priming in the previous experiments, the stimuli were not preceded by primes or pre- and post-masks.

The distribution of ‘Same’ responses is shown in Fig. 6A. Once again, the centroid value was significantly higher for the familiar object (*b* = blur level of 7.93, *SE* = 0.11) than for the novel object (*b* = blur level of 7.43, *SE* = 0.083; Fig. 6B), indicating that the familiar object was perceived as sharper than the novel object at an equivalent level of blur, *t*_24_ = 5.69, *p* < 0.00001, *SE* = 0.087, *Cohen’s d* = 1.138, replicating the results of Experiment 2 without word primes before the displays. As in Experiment 2, the centroids for both *Test* Object types were greater than the value of the *Standard*, indicating the asymmetric application of a ‘Different’ criterion for *Test* objects blurrier versus sharper than the *Standard*. Critically, this occurred when both familiar and novel objects were the *Test* objects and did not prevent the observation of a difference in the centroids for familiar versus novel *Test* objects.

**Figure 6.**
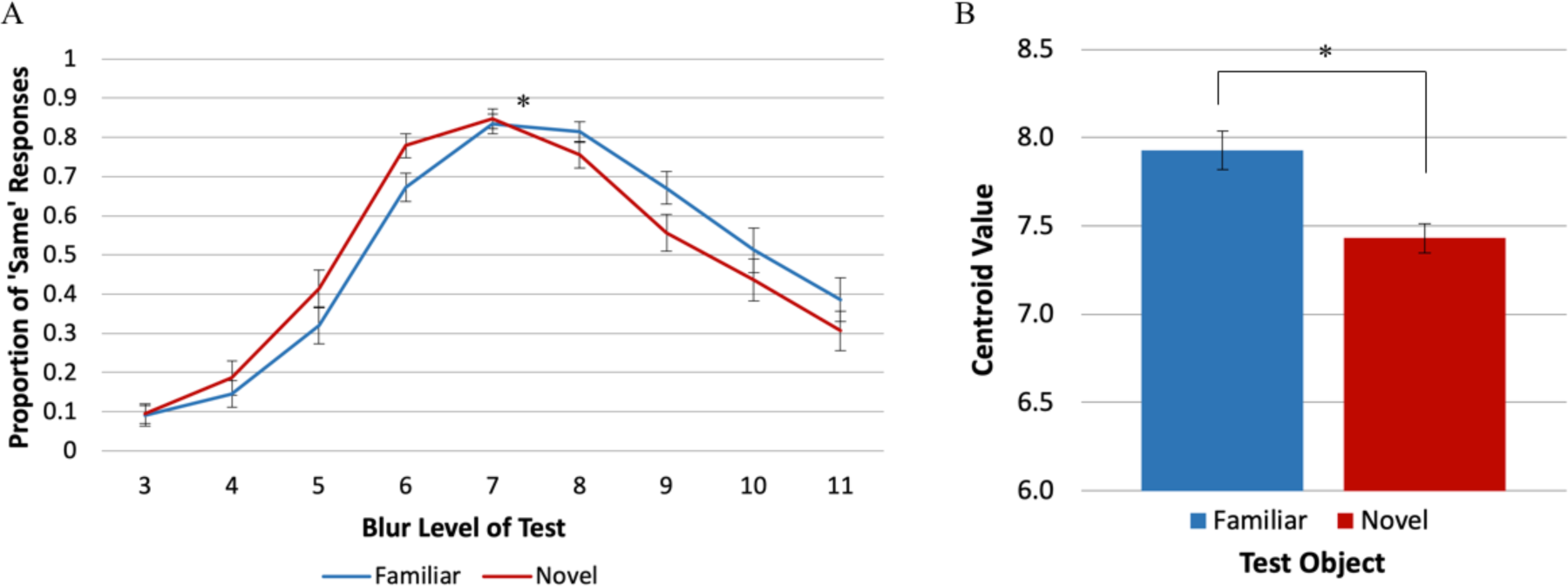
Results of Experiment 3. A) The mean proportion of ‘Same’ responses as a function of the blur level of the *Test* object. B) Centroid value estimates show a main effect of *Test* Object type. For familiar *Test* objects a blurrier stimulus was subjectively equal to a *Standard* novel stimulus of blur level 7. These results, obtained using equality judgments and without primes, are consistent with the hypothesis that for a given level of blur, familiar objects appear sharper than novel objects. Error bars represent Standard Error.

The responses collected in the post-experiment questionnaire revealed that most participants identified the familiar object, with 70% reporting it was a lamp. The most common interpretations for the novel object were ‘Vase’ and ‘Octopus’, with 13% agreement each. Thus, participants were substantially more likely to perceive one of the stimuli as denoting a well-known “familiar” object than the other.

### Experiment 4 – Equality Judgements with Different Stimuli

In Experiment 4, to examine whether this effect generalizes beyond the stimuli used in Experiments 1-3, we assessed equality judgements with two new sets of stimuli: An anchor and a matched part-rearranged novel stimulus (Fig. 1B; Experiment 4A) and a woman and a matched part-rearranged novel stimulus (Fig. 1C; Experiment 4B).

The distribution of ‘Same’ responses in Experiment 4A is shown in Fig. 7A. A shift in the distributions of ‘Same’ responses is evident again, indicating that blurrier borders of familiar *Test* objects were perceived as equal to those of the *Standard* novel object. The centroid value was statistically higher for the familiar object than for the novel object (Fig. 7B; *b*_familiar_ = blur level of 8.28, *SE* = 0.090; *b*_novel_ = blur level of 7.48, *SE* = 0.10), *t*_16_ = 7.06, *p* < 0.00001, *Cohen’s d* = 1.71. These results replicate the effects obtained in the previous experiments where observers reported perceiving the two objects as equal in blur when the familiar *Test* object was at a higher blur level than the novel *Standard* object. The estimated familiarity-dependent sharpening step was 0.40 (*SE* = 0.056) of a blur level. As in Experiments 2-3, the centroid values for both the novel and the familiar *Test* objects were higher than that of the *Standard* (blur level = 7), consistent with an asymmetric application of a ‘Different’ criterion, that we attribute to the nonlinear response to blur^27–28^ affecting the criterion for both types of *Test* objects. In Experiment 4A, 90% of participants identified the familiar object as an anchor correctly. Only one interpretation of the novel object was mentioned by more than one participant: “Yoda” (a “Star Wars” film character), with 15% agreement. Thus, participants were substantially more likely to perceive one of the stimuli as denoting a familiar object than the other.

**Figure 7.**
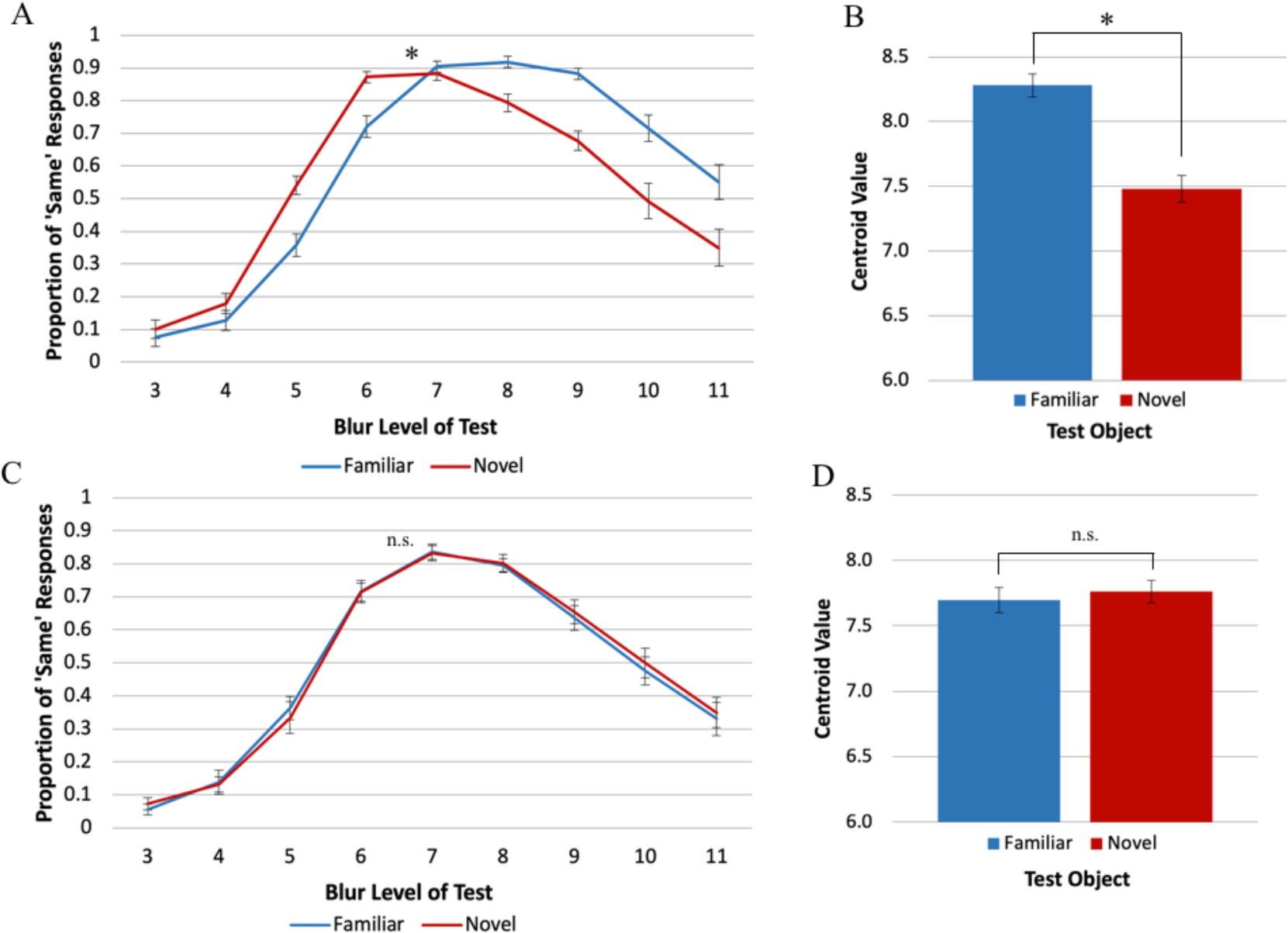
Results of Experiment 4. A&C) The mean proportion of reports that the objects were the same level of blur as a function of the blur level of the *Test* object A) for Exp. 4A and C) for Exp. 4B. B&D) Centroid value estimates for each *Test* Object type B) for Exp. 4A and D) for Exp. 4B. For a *Test* object perceived as more familiar than a novel *Standard* of blur level 7, as in Exp. 4A, a blurrier stimulus is subjectively equal to a *Standard* novel stimulus, consistent with the hypothesis that for a given level of blur, familiar objects appear sharper than novel objects. This result is not observed when both the *Test* object and the Standard object appear familiar, as in Exp. 4B. Error bars represent Standard Error.

In Experiment 4B, we did not obtain the previously-seen shift in perceived blur between the familiar and novel conditions (Fig. 7C). The centroid values obtained for the familiar and the novel objects (Fig. 7D; *b*_familiar_ = blur level of 7.70, *SE* = 0.095; *b*_novel_ = blur level of 7.76, *SE* = 0.085) did not differ statistically, *t*_15_ = −0.83, *p* = 0.42, *Cohen’s d* = 0.21 The responses to the post-experiment questionnaire for Experiment 4B reveal the reason: while 80% of participants were able to correctly name the familiar object as a woman, there was high agreement that the putatively novel object depicted a familiar object: 45% agreed that the novel object resembled a lamp, and 20% agreed it represented a male figure, producing a combined index of 65% familiar object interpretations for the novel stimulus. Thus, Experiment 4B was not a good test of our hypothesis, although it did establish that differences between conditions are not necessarily obtained using this method; differences in perceived familiarity are essential. In Experiment 4B as in Experiments 2-4A, the centroid values for both familiar and novel objects were greater than the *Standard* value of 7, supporting the hypothesis that the asymmetric application of the criterion for ‘Different’ was a consequence of participants’ differential ability to perceive differences in blur across the blur continuum rather than differences in familiarity.

## Discussion

In Experiment 1 participants made comparative judgements, indicating which of two simultaneously displayed blurry objects was ‘Blurrier’; PSEs indicated that for equivalent levels of blur, a familiar object appeared sharper than a novel object, both when it was the *Standard* and when it was the *Test* object. These results were consistent with the hypothesis that object perception in general and object appearance in particular result from the integration of memory representations with the input. For well-known objects, these include many instances in which the objects were fixated and attended outside the laboratory^18^, thus these representations are sharper than the blurry images used in these experiments. The (sharper) activated memory representations of the known object are integrated with the (blurry) sensory input, and this will produce a percept that is sharper than the displayed object. Note that we conceive of object memories as dynamic representations that are updated with each exposure to an object. Nonetheless, experience with novel objects accrued during the experiment cannot produce equivalent results because only blurry versions of the objects were shown. Indeed, we found no evidence that experience with blurry versions of the familiar object during the experiment caused it to appear less sharp in the second half of trials.

Might a response bias interpretation fit our results? Suppose, for instance, that there was a bias to pair “novel” with “blurry;” could it be argued that the results of Experiments 1A and B index response bias rather than appearance? (Note that no such bias is known.) In order to eliminate potential response bias effects, participants in Experiments 2-4 made equality judgements. Anton-Erxleben et al^24^ had replicated previously observed^9^ effects of attention on perceived contrast in experiments measuring equality judgments rather than comparative judgments, thereby ruling out a response bias interpretation for those results^25^. Similarly, our experiments measuring equality judgments replicated the results we obtained with comparative judgments showing that blurry familiar objects appeared sharper than equivalently blurry matched novel objects. Experiment 4B did not show the same pattern of results, but responses to a post-experiment questionnaire revealed that both the familiar and the putatively novel object were perceived as depicting familiar objects. Thus, using the same procedure with two seemingly familiar objects, the difference in perceived sharpness disappeared, confirming that the effects obtained in the previous experiments are indeed mediated by activation of familiar objects memories, and not by a procedural artefact.

In Experiments 2-4 the centroid value for the novel stimulus deviated from the *Standard* in the same direction, albeit to a lesser degree, as that for the familiar stimulus. This pattern was attributed to the asymmetric use of the two criteria for ‘Different’ responses operative in equality judgments, which resulted in a more liberal use of ‘Same’ responses for *Test* objects with larger blur levels than the *Standard*. It is known that sensitivity to differences in blur decreases with increasing blur levels^27, 28^. Perhaps the ‘different because blurrier’ criterion was affected when discrimination was harder. This change in criterion operated equally for familiar and novel *Test* objects, however. Accordingly, the difference between the centroids for the familiar and novel objects can still estimate familiarity-dependent differences in perceived sharpness of object borders. Like Experiment 1, Experiments 2, 3 and 4A showed that blurry familiar objects appear significantly sharper than blurry novel objects.

In Experiments 2-4A, the sharpening step was approximately 1/3 of a blur step; this small effect is likely due to the decreased sensitivity of equality judgements relative to comparative judgements^24–26^. In Experiments 1A and 1B where comparative judgements were used, the average sharpening step was 1/2 a blur step, which is approximately half of the just noticeable difference (JND; approximately one blur level in Experiments 1A and 1B). Despite a reduction in the measured size of the sharpening step in Experiments 1-4A, it remained significantly greater than zero, demonstrating that for a given level of blur, familiar objects appear sharper than novel objects. The magnitude of past experience influences measured in any given experiment likely depends on a variety of factors, including experimental procedure as demonstrated here. Accordingly, we argue that what is critically important is the evidence that familiar objects appear sharper than novel objects, not the degree of sharpening observed in any single experiment. Our results demonstrate that the brain does not produce an exact replica of our surroundings. Instead, via the integration of memory representations from previous experiences, it finds the best interpretation for the often ambiguous input it receives, and this alters object appearance. Here, we have demonstrated that, when objects have blurry borders, well-known (“familiar”) objects appear sharper than novel objects. Future experiments should explore whether other visual properties, such as contrast and spatial frequency are also influenced by prior experience.

Might our results be explained by attention rather than by the integration of incoming information with memories of familiar objects? Some have claimed that attention is automatically allocated to familiar over novel objects^29^. Yet a recent study^30^ showed that previous results taken to support this claim were replicated only under conditions of search involving spatiotemporal uncertainty^31, 32^. Our experiments did not involve search nor was spatiotemporal uncertainty present: each stimulus was shown in one of only two locations at a fixed interval after trial onset; hence it is unlikely that attention was allocated toward the familiar object. Is it possible that when participants found the blur comparison difficult, their uncertainty was high such that familiarity was prioritized for attentional allocation, thereby causing the perceived sharpening of the borders of the familiar objects? Given that the only uncertainty in the equality experiment was whether the two objects were the same or different blur levels, we find this explanation unlikely. Finally, there is some evidence that novel objects are more likely than familiar objects to attract attention^33–35^.

Could claims that attention is attracted by blur explain our results? Both novel and familiar stimuli were presented an equal number of times in all blur levels; hence this explanation cannot account for our data. Moreover, blur detection has been shown to occur pre-attentively^17^. Object memories have also been shown to operate pre-attentively to influence figure assignment^36–42^. Consequently, the explanation in terms of the integration of object memory and current stimulation better fits the data than an explanation in terms of attention.

Previous studies have shown that borders of stimuli such as those tested here can access memory representations of familiar objects very early in processing and serve as priors for figural assignment^4, 5, 36, 38–42^. The present experiments show that the influence of past experiences transcends the effect seen on figural assignment and extends to the subjective appearance of stimuli.

Higher-level influences on the perception of stimuli have also been observed in studies on perceived sharpness of objects in peripheral vision^43–45, cf. 46, 47^. Objects in the periphery tend to appear sharper than an equally-blurry object presented foveally. This “sharpness overconstancy” is thought to arise from a default assumption of sharpness caused by a prevalence of sharp borders in the visual world. Our results may relate to sharpness overconstancy because our stimuli were presented in the near retinal periphery. However, sharpness overconstancy cannot account for the differential sharpening of the familiar relative to the novel objects because both were equidistant from fixation; participants were comparing two peripheral stimuli to each other. Thus, our results add to the literature by showing that this phenomenon varies as a function of the familiarity of the object.

The effects seen in this study may be mediated by feedback from high-level brain regions representing object memories. Recently, the perirhinal cortex (PRC) of the medial temporal lobe (MTL), traditionally considered exclusively a memory-related structure, has been implicated in object perception^48–54^. Barense and colleagues found that the PRC plays a critical role in border familiarity effects on figure assignment^52^ and proposed that the PRC modulates activity in the lower level visual area V2. Subsequent studies reported evidence consistent with this proposal^53–54^; this evidence is consistent with the idea that memory representations of objects are integrated with the input, as we hypothesize here (although we note that there is no evidence that feedback from the PRC in particular mediates the present results).

These results showing that memory and perception interact to affect perceived blur should be of interest to a broad range of scientists who seek to understand the brain (e.g., neuroscientists, computational modelers, psychologists), to computer vision scientists, to designers of autonomous vehicles that must operate in less than ideal conditions, and to those who design visual displays. Elucidating the contributions of higher-order regions of the brain to visual perception brings us closer to understanding the intricate processes that give rise to the perceptions that guide our behaviour and allow us to interact with our environment.

## Methods

The methods used in these experiments were carried out in accordance with guidelines established by, and the explicit approval of, the Human Subjects Protection Program at the University of Arizona. The experiments were conducted on a Dell OptiPlex 9020 computer and displayed on a 24-in. AOC monitor at 1920 x 1080 resolution and a 100Hz refresh rate. Participants used a standard keyboard to navigate from trial to trial. In Experiments 1 and 2, the keyboard was used to make responses. In Experiments 3-5, participants used a separate number pad to make responses. The experiment was programmed using MATLAB (The Mathworks, Natick, MA) R2015a and the Psychophysics Toolbox-3.0.12^55–56^.

The stimuli were black silhouettes (luminance 0.1 ftL) of familiar and novel objects, presented on a medium grey background (luminance 31.7 ftL). In Experiments 1-3, the familiar stimulus depicted a table lamp (Fig. 1A); in Experiments 4A and 4B the familiar stimuli depicted an anchor (Fig. 1B) and a standing woman (Fig. 1C). To control for differences in low-level features, the novel stimulus was constructed by spatially rearranging the parts of the familiar stimulus, with parts defined as lying between two successive minima of curvature ^cf. 57–58^. The number of black pixels in the paired familiar and novel stimuli was equal.

In each of 864 trials, a familiar and a novel stimulus were shown to the right and left of fixation. Before each trial, each stimulus was blurred to its assigned blur level using MATLAB’s 2-D Gaussian filtering function *imgaussfilt.* The two inputs for this function were the unblurred stimulus image and a value specifying the standard deviation of the isotropic Gaussian smoothing kernel centred on each pixel that blurs surrounding pixels. The size of the filter applied to the image pixels was determined by the default equation:

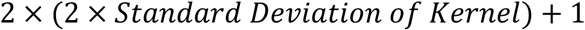

This resulted in the filter size increasing by 4 pixels for every one unit increase in standard deviation. Pixel size was approximately 0.01586° of visual angle. We varied the degree of blurriness of the stimuli by varying the value of the standard deviation from 3 (least blurry) to 11 (blurriest) in intervals of 1. Unblurred stimuli were never shown in this experiment. Here, we refer to the standard deviation of the smoothing kernel as the “blur level” of the stimulus. The filter size was 13×13 pixels when blur level = 3 and 45×45 pixels when blur level = 1.

In each trial, one stimulus was the *Standard* and the other was the *Test*; each served as *Standard* and *Test* equally often. The *Standard* was held constant at blur level = 7. The *Test* object varied from blur levels of 3 to 11. The number of black pixels in the familiar and novel objects did not differ within each of the three stimulus pairs that were used. The stimuli were centred 4.13° from fixation. Participants viewed the displays from a distance of 98.5 cm. When unblurred, the lamp stimulus pair subtended 2.67° H X 2.04° W of visual angle, the anchor the stimulus pair subtended 2.85° H X 2.56° W of visual angle, and the woman stimulus pair subtended 2.62° H X .87° W of visual angle.

We examined whether we manipulated familiarity effectively with a post-experiment questionnaire where participants were asked to name the two objects that were displayed during the experiment. The answers to this question are summarized in the Results sections. Additionally, while this experiment was underway an online Amazon Turk experiment was conducted to determine the familiarity and novelty of a larger set of stimuli^59^. Thirty-two participants viewed half portions of each stimulus (see Fig. 8), and listed the familiar objects resembled by each stimulus. For the stimuli used in our experiments the results obtained in the online experiment corroborate the results we obtained from the participants in our experiments.

**Figure 8.**
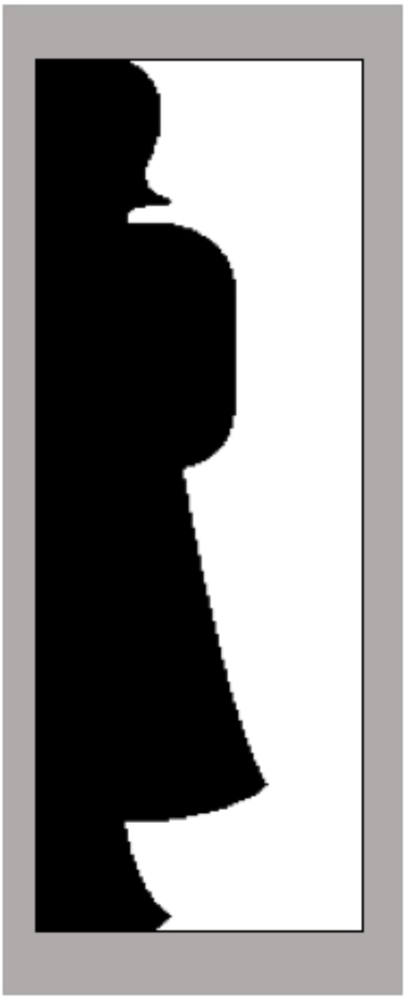
Example of bipartite display. Bipartite displays were used in an online experiment to assess familiarity/novelty of stimuli. In this case, the black region delineates the silhouette of a standing woman.

In Experiments 1A, 1B, and 2, the stimuli were preceded by one of three types of primes: (1) *identity primes*, intended to induce an expectation for the named stimulus, (2) *unrelated* primes (matched for word frequency to the identity primes^60^), intended to induce an expectation for an unrelated object from a different superordinate category, and (3) *control* primes, where no word was presented prior to stimulus presentation. All primes were matched in word length. For those on which the familiar object was the *Test*, the identity prime was “lamp”, the unrelated prime was “hawk”, and the control prime was “xxxx.” For trials on which the novel object was the *Test*, the primes could not be classified as related or unrelated. Accordingly, we chose “dummy” primes from the same categories as the primes for the familiar object. For the novel objects, the dummy primes for the *identity, unrelated*, and *control* conditions were: “pole”, “chin”, and “xxxx.” All Prime types were presented equally often. The prime words “lamp” and “pole” subtended 0.87° X 2.44°, “hawk” subtended 0.64° X 2.50°, “chin” subtended 0.64° X 2.44° and “xxxx” subtended 0.47° X 2.44° of visual angle. The size of the monitor was 29.8cm H X 52.5cm W; it subtended 17. 2° H X 29.8° W of visual angle at the viewing distance used.

Word primes were both pre- and post-masked to render them unconscious. In Experiment 1A, the pre-mask and post-mask were unique strings of random consonants. In Experiments 1B and 2, the pre-mask was changed to a string of #’s. The masks were the same length as the prime words.

### Experiments 1A and 1B – Comparative Judgements

Before beginning the experiment, participants viewed the instructions on the computer screen while the experimenter read them aloud. The instructions explained that they would see two objects on each trial, one to the left and one to the right of a fixation cross. Their task was to report which object was blurrier. They were shown examples of the two stimuli and instructed to make their responses with a button press using the left and right arrow keys. They were told they would see the same two objects throughout the experiment. Next, they were guided through a typical trial sequence (Fig. 2). They then completed 3 blocks of 10 practice trials each. Prime type and blur level of the *Test* objects (3 - 11) were balanced in these practice trials. Each stimulus appeared as *Standard* and as *Test* and on each side of the screen an equal number of times. Each block of practice trials was presented in a random order so that each participant completed a different sequence of practice trials. Three practice blocks accustomed participants to viewing short exposure durations: the post-mask for the prime and the experimental stimuli were shown for increasingly shorter exposure durations across the three blocks (twice as long in the first practice block as on experimental trials, 1.5*X* as long in the second practice block, and 1.0*X* as long in the third practice block).

Participants completed 864 experimental trials. They were instructed to fixate on a cross in the centre of the screen throughout the entire trial. They initiated each trial by pressing the space bar. In Experiment 1A, a pre-mask appeared for 350 ms, followed by the prime (40 ms), followed by the post mask (80 ms). In Experiment 1B the pre-mask was a string of #’s instead of a random consonant sequence, the word prime was lengthened to 50 ms, and the post-mask was shortened to 70 ms. After the post mask, the stimuli were displayed for 180 ms, a duration too short for a saccade to one side or the other. The familiar and novel stimuli occurred equally often on the left and the right of fixation. In each trial, one of the stimuli was the *Standard* (blur level = 7); the other, *Test*, stimulus was presented at blur levels ranging from 3 - 11 equally often. The familiar stimulus was the *Test* on half the trials; the novel stimulus was the *Test* on the other half the trials. There were 16 trials for each combination of stimulus condition (familiar vs novel *Test* stimulus), priming condition (identity, unrelated, control), and *Test* stimulus blur level (3-11). All trial types were randomly intermixed.

After the stimuli disappeared, participants had up to 3 seconds to report whether the object on the right or on the left was ‘Blurrier.’ Participants were instructed to report which object was blurrier, as opposed to which was sharper in order to reduce the possibility that any bias to pair the familiar object with the instructed response would favour our hypothesis^22^.

Participants responded using their right hand on the left and right arrow keys, where they pressed the left arrow key for a left response, and the right arrow key for a right response.

#### Statistical Analysis

Trials on which a response was not recorded during the 3s response window were excluded. Trials with incorrect display timing were also removed (<1% of trials). This was determined by a MATLAB script written to verify the display timing for each frame in the experiment. For each participant, the data were separated by Prime type (identity, unrelated, control) and *Test* Object type (familiar, novel). For each of these conditions, the number of times the *Test* object was chosen as ‘Blurrier’ was averaged for each blur level. These averages served as input for the curve-fitting toolbox in MATLAB. Using a Non-Linear Least Squares method, the distribution of each participant’s responses was fit to a Logistic Function,

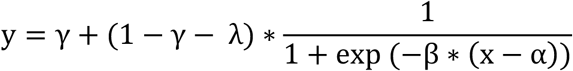

where λ is the lapse rate, γ is the guess rate, α is the threshold parameter, and β is the slope parameter. The threshold value corresponds to the location of the logistic curve where participants are at chance level reporting either side as the blurrier option, indicating the point of subjective equality (PSE). We obtained the threshold parameter value for each participant’s distribution, along with a confidence interval for this value. An outlier removal procedure consisted of removing any participant whose confidence interval for the threshold parameter was lower or higher than two standard deviations from the mean confidence interval for all participants. Three rounds of this procedure were performed. The PSE values were submitted to a 2 × 3 repeated measures ANOVA with *Test* Object type (familiar, novel) and Prime type (identity, unrelated, control) as factors.

### Experiments 2-4 Equality Judgements

Participants were instructed to report whether the two objects shown on each trial were the same or different levels of blur. They recorded their responses using a separate number pad rotated 90° to the right, with keys on only the numbers 1 (at the top) and 3 (at the bottom). Participants were instructed to press the top key if the two stimuli were equal in blur or the bottom key if they were different. In other respects, the Experiment 2 procedure was the same as that of Experiment 1B. The procedure for Experiment 3 was the same as Experiment 2, except that no word primes were presented at any point during the experiment. Experiments 4A and 4B were run interleaved in that when participants entered the laboratory, they were assigned to one of the two experiments using an ABBA procedure. Participants took part in one experiment only. The procedure was the same as in Experiment 3, with the exception of the stimuli. In Experiment 4A the familiar stimulus was the silhouette of an anchor and in Experiment 4B the familiar stimulus was the silhouette of a woman.

#### Statistical Analysis

For each participant, the data were separated by Prime type (identity, unrelated, control) and *Test* Object type (familiar novel), in a manner similar to Experiments 1A and 1B. The distribution of each participant’s responses was fit to a normal Gaussian distribution using a Non-Linear Least Squares method,

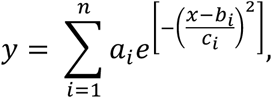

where *a* is the amplitude, *b* is the centroid value, *c* is related to the width of the peak, and *n* is the number of peaks to fit (in this case, one peak). The centroid value, *b*, corresponds to the location of the peak, where the probability of a ‘Same’ response is highest. A two-way repeated measures ANOVA was performed on the results of Experiment 2 to assess the effect of Prime type and *Test* Object type on the centroid value. A paired-sample, two-tailed t-test was used to compare the centroid values for the familiar and novel *Test* Object types in Experiments 3 and 4.

#### Participants

All participants were University of Arizona (UA) undergraduate students (ages 17-23) who participated to fulfil a class requirement. Participants and/or their legal guardian gave informed consent on a form approved by the UA Human Subjects Protection Program. Participants who were 18 years of age or older completed this form before taking part in the experiments. Participants who were 17 years old were given the option to participate in order to fulfill their class requirement and to have a consent form mailed to their legal guardian. Their data were analysed only after we received a signed consent form in the mail. Participants had normal or corrected to normal vision (at least 20/30) assessed by responses to a Snellen Eye Chart Test. Before beginning data collection, two discard criteria were established for all experiments in this article: Criterion 1 was “failure to follow task instructions” and criterion 2 was “pressing the wrong button in 30% of trials or more” as reported in the post-experiment questionnaire.

Experiment 1A: The data of 14 participants (7 male, 7 female) are reported. Two other participants were discarded based on criterion 1; eight others were excluded after the outlier removal procedure. Experiment 1B: The data of 14 participants (5 male, 9 female) are reported. Two other participants were discarded based on discard criterion 1; eight others were excluded after the outlier removal procedure. Experiment 2: The data of 23 participants (11 male, 12 female) are reported. Six other participants were excluded after the outlier removal procedure; none were discarded based on discard criteria. Experiment 3: The data of 25 participants (12 male, 13 female) are reported. Two other participants were discarded based on discard criterion 1; four were discarded based on criterion 2; and five others were excluded after the outlier removal procedure. Experiment 4A: The data of 17 participants (10 male, 7 female) are reported. Four other participants were discarded based on discard criterion 1; three others were excluded after the outlier removal procedure. Experiment 4B: The data of 16 participants (9 male, 7 female) are reported. Four other participants were discarded based on discard criterion 1; one was discarded based on criterion 2; and four others were excluded after the outlier removal procedure.

## Data Availability

All data that support these findings are publicly available at https://doi.org/10.5281/zenodo.3325023

## Code Availability

The code necessary to reproduce these findings and analyses is available at https://github.com/dianaperez25/BlurPerception

## Acknowledgements

This research was funded by a grant from the Office of Naval Research (N00014-14-1-067). The funders had no role in study design, data collection and analysis, decision to publish or preparation of the manuscript. The authors thank R. Skocypec, C. Flowers, and M. Jernigan for helpful discussion and feedback during this project, E. Wolf and E. Idowu for help collecting data, and to the Undergraduate Biology Research Program (UBRP) for their continuous support. We would especially like to thank our reviewer, M. Georgeson, for his careful review and enormously helpful comments.

## Author Contributions Statement

S.M.C. and M.A.P. conceived the study. S.M.C., D.C.P. and M.A.P. designed the experiments.

S.M.C. and D.C.P. collected and analysed the data. D.C.P. wrote the first draft of the manuscript.

M.A.P. provided direct supervision and feedback during study design, data collection, data analysis and interpretation, and manuscript preparation stages. All authors approved the final version of the manuscript.

